# Increased Postprandial Metabolic Flexibility is Associated with Higher Body Fat Percentages in Healthy Young Adults

**DOI:** 10.1101/2024.08.23.609405

**Authors:** Nicholas A. Foreman, Sahil Rajwade, Jaiden Bluth, Lauren C. Skoglund, Audrey M. Letts, Loretta DiPietro, Adam Ciarleglio, Matthew D. Barberio

## Abstract

**PURPOSE:** Because higher adiposity is associated with cardiometabolic disease, we assessed the relationship between body composition (body fat percentage; BF%) and postprandial metabolic flexibility.

**METHODS:** Young adults (n = 27, n = 15 females, BMI = 27.1 ± 4.5; BF% = 30.4 ± 8.7) without overt pathology completed a 100g oral glucose tolerance test (OGTT). Indirect calorimetry before (fasting) and following (30, 60, 90, 120 min) consumption was used to calculate respiratory exchange ratio (RER) and oxidation of carbohydrates (CHO) and fats (FOX). Serum and plasma were collected at corresponding time points and analyzed for blood glucose, insulin, and NEFAs. Data from individuals with normal weight were compared to those with overweight/obesity by two-way repeated measures ANVOA. The effect of BF% on postprandial metabolism was tested via linear mixed models while adjusting for potential confounders.

**RESULTS:** During the OGTT, blood glucose, serum insulin, plasma lactate, RER, and CHO all significantly increased while plasma NEFAs and whole-body FOX decreased (all p<0.05). BF% modified the relationship between postprandial RER and time (p = 0.019) as well as postprandial CHO and time (p = 0.023) without an effect of BF% on FOX; individuals with higher BF% increase their RER and CHO faster and to a greater extent than those with lower BF%.

**CONCLUSION:** Body fat percentage is associated with greater postprandial metabolic flexibility during an OGTT in young adults. Despite increased adiposity, metabolic flexibility may be preserved, representing a compensatory adaptation to decreased glucose storage in the postprandial period.

## Introduction

Metabolic flexibility is described as the appropriate shift in substrate oxidation based upon substrate availability (e.g. fasting to fed) or metabolic demand (e.g. exercise). It was originally described as the change in the respiratory quotient (RQ) across the leg limb from rest to insulin stimulation during the euglycemic-hyper-insulinemic clamp (EHC).^1, 2^ In the two decades since the concept of metabolic flexibility was described in humans, several methodological approaches to assess metabolic flexibility have been utilized. These include indirect calorimetry during the EHC, during oral glucose tolerance tests (OGTT), and during mixed-meal feeding challenges (MMC).^1, 3^ Individuals with overt metabolic pathologies, such as type 2 diabetes and insulin resistance, as well as individuals with obesity based on BMI, have been described as metabolically inflexible.^1^ That is, these individuals showed blunted (reduced) changes in the respiratory exchange ratio (RER) during metabolic challenges such as EHC and feeding.

In the absence of overt metabolic pathology or metabolic dysfunction, transitioning from the fasted to the fed state involves shifting away from aerobic fatty acid oxidation to aerobic glucose oxidation and, to a lesser degree, glycolytic energy production.^4^ This shift results in an anabolic environment, in which exogenous energy is stored, as compared to the catabolic environment of fasting. Elevated insulin, which is a primary driver of the metabolic shift following feeding, suppresses adipose tissue lipolysis and promotes the uptake, storage, and oxidation of exogenous macronutrients.^4^ Adipose tissue serves a critical role in buffering fatty acid flux in the circulation during this postprandial period; it is also a site of significant glucose uptake for lipogenesis and has been identified as a central regulator of metabolic flexibility.^2, 4^

The presence of obesity and increased fat mass is believed to negatively impact metabolic flexibility.^5, 6^ Individuals with obesity have higher resting RER indicating a preference for higher carbohydrate oxidation in the fasting state. In obesity, adipose tissue lipolysis is also more resistant to insulin-induced suppression in feeding. However, studies have not thoroughly analyzed the effect of adiposity on postprandial metabolic flexibility. By dividing participants into high vs. low metabolic flexibility, Sparks et al. (2009) demonstrated that healthy young males with low metabolic flexibility during insulin stimulation (EHC) had significantly higher body fat percentage and larger fat cell size.^6^ Follow-up analysis in this cohort demonstrated that females were more metabolically flexible than their male counterparts despite significantly higher body fat percentages.^5^ Conversely, individuals classified as having high metabolic flexibility to an OGTT had similar body fat percentages but a larger subcutaneous adipose tissue (SAT) as compared to those described as having low metabolic flexibility.^7^ This discrepancy was highlighted in a recent systematic review by Glaves et al. (2021), who found inconsistent and unclear relationships between adipose tissue volume, function, and characteristics in studies using EHC, OGTTs, and MMCs.^3^

The primary objective of this study was to thoroughly assess the relationship between adiposity and postprandial metabolic flexibility during an OGTT. We recruited 27 healthy male and female young adults (18 - 40 years old) with body mass index (BMI) between 18.5-39.9. Using indirect calorimetry, we assessed postprandial metabolic flexibility (RER, Carbohydrate Oxidation [CHO], and Fat Oxidation [FOX]) during an OGTT. We hypothesized that individuals with higher body fat percentages would demonstrate significantly reduced postprandial metabolic flexibility (RER and CHO) throughout the OGTT.

## METHODS

### Participants

We performed a cross-sectional, experimental study from January 2023 to January 2024. Potential participants were recruited and screened based on sex, body mass index (BMI) classification, and lack of known clinical pathology from the George Washington University campus and surrounding metropolitan area. Participants were required to be between 18 and 40 years old, have a BMI between 18.5 and 39.9, be free of clinically diagnosed cardiometabolic disease, and not be on prescription medications for hypertension, blood lipids, or blood glucose homeostasis. Participants who were pregnant or taking medications for cardiometabolic disease were excluded. Recruitment was stratified by BMI classification to yield a representative sample across the BMI continuum. In response to flyers and emails, fifty-two participants were screened for participation. Of these, thirty-four were enrolled in this study; seven participants did not complete the study (3 due to scheduling, 3 due to loss to follow-up, 1 due to equipment malfunction), giving complete data for 27 participants. Participant characteristics are reported in Table 1. All study procedures were approved by the George Washington University IRB (IRB #202737). Prior to enrollment, participants read and signed an IRB-approved informed consent, and all study procedures were conducted in accordance with the Declaration of Helsinki.

**Table 1:**
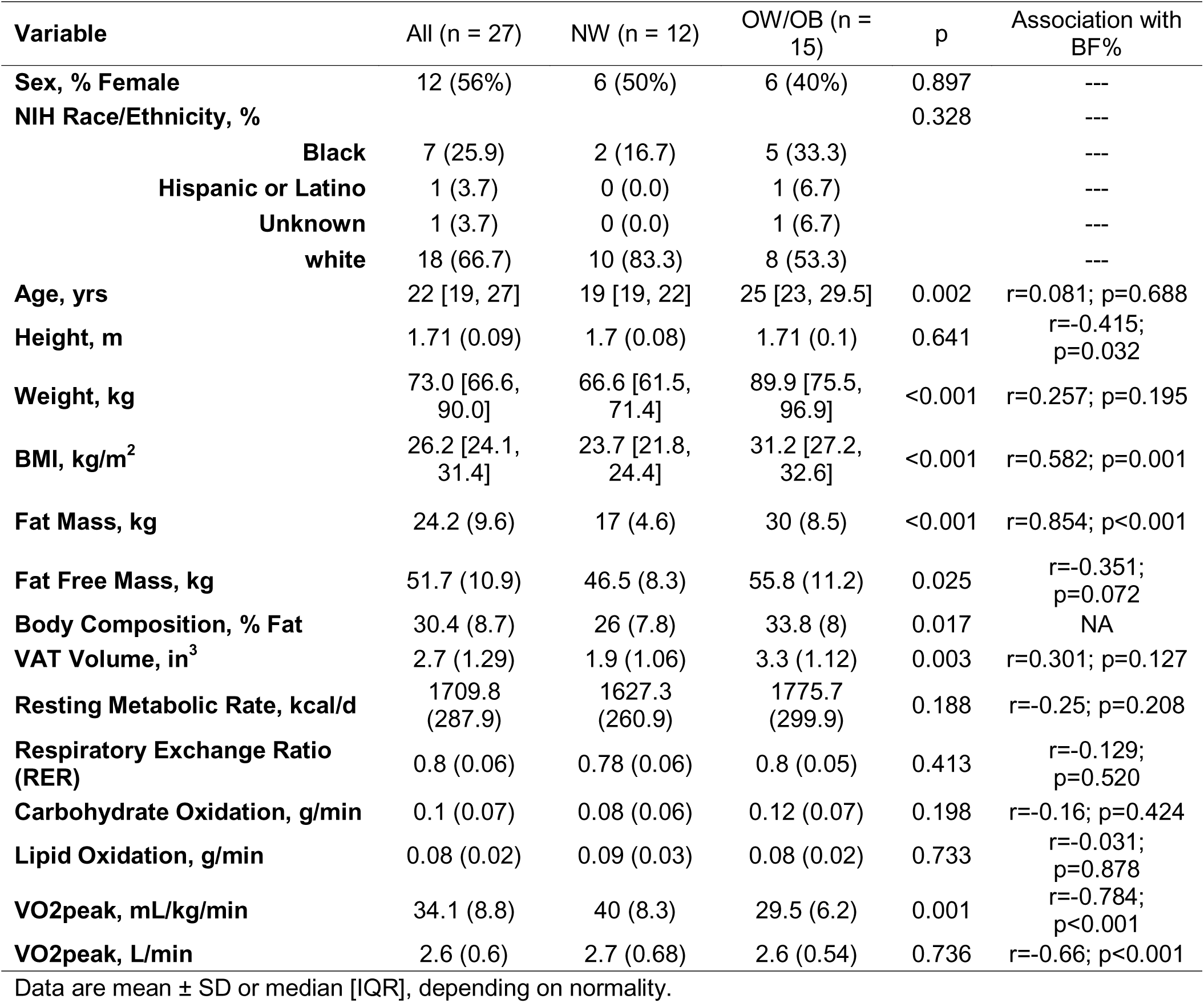
Participant Characteristics.

| Variable | All (n = 27) | NW (n = 12) | OW/OB (n = 15) | p | Association with BF% |
| --- | --- | --- | --- | --- | --- |
| **Sex, % Female** | 12 (56%) | 6 (50%) | 6 (40%) | 0.897 | --- |
| **NIH Race/Ethnicity, %** |  |  |  | 0.328 | --- |
| **Black** | 7 (25.9) | 2 (16.7) | 5 (33.3) |  | --- |
| **Hispanic or Latino** | 1 (3.7) | 0 (0.0) | 1 (6.7) |  | --- |
| **Unknown** | 1 (3.7) | 0 (0.0) | 1 (6.7) |  | --- |
| **white** | 18 (66.7) | 10 (83.3) | 8 (53.3) |  | --- |
| **Age, yrs** | 22 [19, 27] | 19 [19, 22] | 25 [23, 29.5] | 0.002 | r=0.081; p=0.688 |
| **Height, m** | 1.71 (0.09) | 1.7 (0.08) | 1.71 (0.1) | 0.641 | r=-0.415; p=0.032 |
| **Weight, kg** | 73.0 [66.6, 90.0] | 66.6 [61.5, 71.4] | 89.9 [75.5, 96.9] | <0.001 | r=0.257; p=0.195 |
| **BMI, kg/m<sup>2</sup>** | 26.2 [24.1, 31.4] | 23.7 [21.8, 24.4] | 31.2 [27.2, 32.6] | <0.001 | r=0.582; p=0.001 |
| **Fat Mass, kg** | 24.2 (9.6) | 17 (4.6) | 30 (8.5) | <0.001 | r=0.854; p<0.001 |
| **Fat Free Mass, kg** | 51.7 (10.9) | 46.5 (8.3) | 55.8 (11.2) | 0.025 | r=-0.351; p=0.072 |
| **Body Composition, % Fat** | 30.4 (8.7) | 26 (7.8) | 33.8 (8) | 0.017 | NA |
| **VAT Volume, in<sup>3</sup>** | 2.7 (1.29) | 1.9 (1.06) | 3.3 (1.12) | 0.003 | r=0.301; p=0.127 |
| **Resting Metabolic Rate, kcal/d** | 1709.8 (287.9) | 1627.3 (260.9) | 1775.7 (299.9) | 0.188 | r=-0.25; p=0.208 |
| **Respiratory Exchange Ratio (RER)** | 0.8 (0.06) | 0.78 (0.06) | 0.8 (0.05) | 0.413 | r=-0.129; p=0.520 |
| **Carbohydrate Oxidation, g/min** | 0.1 (0.07) | 0.08 (0.06) | 0.12 (0.07) | 0.198 | r=-0.16; p=0.424 |
| **Lipid Oxidation, g/min** | 0.08 (0.02) | 0.09 (0.03) | 0.08 (0.02) | 0.733 | r=-0.031; p=0.878 |
| **VO<sub>2</sub>peak, mL/kg/min** | 34.1 (8.8) | 40 (8.3) | 29.5 (6.2) | 0.001 | r=-0.784; p<0.001 |
| **VO<sub>2</sub>peak, L/min** | 2.6 (0.6) | 2.7 (0.68) | 2.6 (0.54) | 0.736 | r=-0.66; p<0.001 |

Data are mean ± SD or median [IQR], depending on normality.

### Experimental Design

Participants reported to the Metabolic and Exercise Testing Laboratories in the morning between 0800 and 0900 on two separate days. Each visit was separated by at least 48 hours. At the first visit, participants completed written questionnaires for medical and physical activity history, assessment of body composition, and cardiorespiratory fitness. Participants returned to the lab for assessment of postprandial substrate metabolism following an OGTT.

For each visit, participants were instructed to arrive after 10 hours of fasting and to abstain from caffeine consumption for 12 hours and from any recreational drug use or structured exercise for 24 hours. Upon arrival, hydration was measured using a portable refractometer. If urine specific gravity was above 1.02, participants were given eight ounces of water. Body weight was measured using a mechanical column scale (Seca 700; Hamburg, Germany).

### Body mass and body composition

Height was measured using a wall-mounted stadiometer (Seca 216; Hamburg, Germany) to the nearest 0.1 centimeters. Weight was measured using the digital scale as part of the bioelectrical impedance analysis (InBody 770; Seoul, South Korea). Dual X-ray absorptiometry (GE Lunar iDXA; GE Healthcare, Chicago, IL, USA) with manufacturer software (enCORE Forma version 15) was used to measure body fat percentage, fat-free mass (FFM), fat mass, and visceral adipose tissue (VAT) volume. Before the measurement, participants removed all jewelry and were instructed to wear clothing free of metal zippers or reflective surfaces.

### Exercise Testing

Following body composition assessment, participants completed a graded exercise test on a cycle ergometer (Monark LC6 novo; Varberg, Sweden) to volitional exhaustion. During cycling, expired gases were measured using a breath-by-breath metabolic cart (Cosmed Quark CPET; Rome, Italy), and heart rate was monitored using a chest strap (Polar H10; Kempele, Finland). All participants completed three 180-second stages of 25, 50, and 75W each before resistance increased by 25W per minute. The test continued until the participant was unable to continue cycling or the investigator observed a plateau in oxygen consumption (VO_2_). Breath-by-breath data were averaged into five-second bins. Because not every participant satisfied the criteria for VO2max,^8^ the highest 15-second value is reported as VO2peak.

### Oral Glucose Tolerance Test with Indirect Calorimetry and Blood Sampling

On a separate day, participants arrived following the same pre-testing guidelines outlined above. Assessment of body weight and hydration were performed, and the participant rested in a reclined chair in a dimly lit room for at least 10-15 minutes. A venous catheter was placed in an antecubital vein and fasting whole blood was sampled for plasma (K-EDTA) and serum (Serum Separator Tube); saline was used to keep the catheter patent between blood draws. After the resting blood draw, indirect calorimetry was conducted via the canopy method (Parvo Medics TrueOne 2400; Salt Lake City, Utah) while the participant lay supine on a padded table for determination of fasting metabolic measures. Following this, subjects consumed a 100g glucose beverage (GlucoCrush Cardinal Health; Dublin, OH) and were instructed to finish consumption of the drink within 5 minutes. After finishing the drink, subjects returned to the supine position for the remainder of the test. Indirect calorimetry was used to assess postprandial substrate oxidation at the following postprandial intervals (min): 15 – 30, 45 – 60, 75 – 90, and 105 – 120.

### Determination of substrate oxidation and metabolic rate

For each 15-minute indirect calorimetry measurement, expired gases were collected from a mixing chamber at 5-second intervals, and the final 8 minutes were used for analysis. Respiratory exchange ratio (RER) was calculated as the ratio between exhaled carbon dioxide (VCO_2_) and VO_2_. Metabolic rate and rates of carbohydrate (CHO) and fat oxidation (FOX) were calculated using published equations for fat oxidation as [VO_2_ (L/min) x 1.67) – (VCO_2_ (L/min) x 1.67)] and carbohydrate oxidation as [VCO_2_ (L/min) x 4.55) – (VO_2_ (L/min) x 3.21)].^9^ For metabolic rate calculations, urinary nitrogen excretion was assumed to be zero.

### Laboratory measurements and calculations

Fasting and postprandial blood glucose were determined in fresh whole blood (OneTouch Ultra, Lifescan, Inc.; Milpitas, CA). Fasting and postprandial serum samples were used to measure insulin (Insulin Human ELISA; Invitrogen Inc. CA), and non-esterified fatty acids (NEFA; LabAssay NEFA; FujiFilm; Richmond, VA). Fasting serum samples were used to determine C-reactive protein (CRP; CRP Human ELISA; Invitrogen Inc., CA), leptin (Leptin Human ELISA; Invitrogen Inc., CA), and adiponectin (Adiponectin human ELISA; Adipogen Life Science; San Diego, CA). Manufacturer instructions were followed for conducting and analyzing all assays. Manufacturer internal controls were assessed along with coefficients of variations for biological replicates. Samples whose performance was assessed to be outside of manufacturer recommendations were re-analyzed. Fresh whole blood (fasting) was used to determine HbA1C% (PTS Diagnostics; Whitestown, IN) and blood lipid profiles (CardioChek; PTS Diagnostics; Whitestown, IN) via point-of-care analyzers.

Matsuda Index was calculated as follows: 10,000/√ [fasting glucose (mg/dl) × fasting insulin (µIU/mL)] × [mean glucose (mg/dl) × mean insulin (µIU/mL) during the OGTT before normalizing to the 100g dose.^10^ Adipose tissue insulin resistance (Adip-IR) was calculated using fasting values as fasting insulin (µIU/mL) * fasting NEFA (mEq/L) and using postprandial OGTT values (30, 60, 90, 120) as mean plasma NEFA (mEq/L) x mean serum insulin (µIU/mL).^11^ Fasting insulin and glucose measurements were used to calculate the homeostatic model assessment of insulin resistance (HOMA-IR) as [fasting insulin (µIU/mL) - (fasting glucose (mg/dl)/18.156)] / 22.5.^12^

### Statistical Analyses

We report data as mean (SD) or median (interquartile range, IQR) both overall and by BMI status for participant characteristics (**Table 1**) and clinical measures (**Table 2**). We compared race categories between the sexes using Fisher’s exact test and report them as frequency (percent). We computed and tested the association between each of the clinical variables and percent body fat (BF%) using Pearson correlation coefficients. To describe the change in each postprandial outcome variables during the OGTT (i.e., RER, CHO, FOX, glucose, insulin, lactate, and NEFA) between individuals with normal weight (NW) and individuals with overweight and obesity (OW/OB) as defined by the BMI, we used two-way, repeated measures analysis of variance (ANOVA) with time modeled as a categorical variable. Contrasts were set *a priori* as the comparison from baseline to each postprandial timepoint (i.e., 0 vs. 30, 0 vs. 60, etc.). Analysis and figures were conducted in OriginPro 2020.

**Table 2:** Clinical Data.

| Variable | All (n = 27) | NW (n = 12) | OWOB (n = 15) | p-value | Association with BF% |
| --- | --- | --- | --- | --- | --- |
| Blood Glucose (mg/dL) | 100 (8) | 98 (12) | 101 (5) | 0.315 | r=0.036; p=0.858 |
| Serum Insulin (microIU/mL) | 14.2 (8.1) | 11.7 (5.8) | 16.3 (9.4) | 0.17 | r=0.404; p=0.050 |
| NEFA (mEq/L) | 0.42 [0.28, 0.49] | 0.34 [0.25, 0.47] | 0.43 [0.32, 0.49] | 0.344 | r=0.339; p=0.098 |
| Serum Lactate (mM) | 1.58 (0.42) | 1.43 (0.33) | 1.72 (0.45) | 0.09 | r=0.237; p=0.266 |
| HbA1c (%) | 5.1 [4.8, 5.25] | 5.05 [4.8, 5.12] | 5.2 [4.85, 5.4] | 0.154 | r=-0.221; p=0.269 |
| Total Cholesterol (mg/dL) | 160.4 (33.9) | 139 (22.8) | 175.6 (32.7) | 0.006 | r=0.198; p=0.353 |
| HDL (mg/dL) | 51.0 [47.8, 59.5] | 52.0 [46.2, 64.8] | 50.5 [48.2, 56.5] | 0.769 | r=0.086; p=0.691 |
| LDL (mg/dL) | 92.3 (29.6) | 71.1 (20.8) | 102.9 (28) | 0.016 | r=0.085; p=0.715 |
| Triglycerides (mg/dL) | 74.5 [66.2, 93.0] | 73.0 [44.5, 76.0] | 82.0 [67.8, 96.8] | 0.135 | r=-0.016; p=0.940 |
| Matsuda Index | 4.24 [2.58, 6.00] | 5.35 [4.03, 7.48] | 3.06 [2.46, 4.78] | 0.082 | r=-0.616; p=0.001 |
| HOMA-IR | 3.1 [2.2, 4.3] | 2.5 [1.9, 3.5] | 3.7 [2.9, 4.6] | 0.207 | r=0.398; p=0.054 |
| AdipIR (OGTT) | 7.7 [4.4, 12.2] | 4.2 [3.0, 10.1] | 8.9 [6.7, 30.4] | 0.035 | r=0.481; p=0.017 |
| AdipIR (fasting) | 5.2 [3.0, 6.9] | 3.5 [2.4, 5.9] | 5.3 [4.1, 9.3] | 0.167 | r=0.443; p=0.030 |
| Adiponectin (ng/mL) | 4613 [3955, 6013] | 4613 [4048, 6874] | 4540 [3307, 5426] | 0.467 | r=-0.151; p=0.470 |
| Leptin (pg/mL) | 118.1 [53.8, 148.5] | 89.8 [36.9, 124.1] | 134.0 [84.3, 198.2] | 0.085 | r=0.817; p<0.001 |
| CRP (mg/L) | 0.48 [0.26, 1.2] | 0.48 [0.23, 1.1] | 0.49 [0.29, 1.1] | 0.727 | r=0.091; p=0.664 |

Data are mean ± SD or median [IQR], depending on normality. VAT volume was adjusted with a cube root transformation. Insulin and NEFA concentrations were log transformed.

To determine the effect of body composition on postprandial metabolism, linear mixed-effects models were used. For each postprandial outcome variable, we included fixed effects for time as a continuous variable, percent body fat (BF%), time x BF% interaction, sex, age, Matsuda index, AdipIR, visceral adipose tissue volume, and the fasted outcome measure. A subject-specific random intercept term was included to account for repeated measures. To account for the possible nonlinear effect of time on postprandial metabolism (i.e., a nonlinear change in metabolism), models with an additional quadratic time term and quadratic time x BF% were considered. Models with and without the quadratic time terms were compared using likelihood ratio tests to determine if the quadratic terms should be retained. Variables in the models were tested for normality with the Shapiro-Wilk test and transformed. As a result, insulin and NEFA concentrations were transformed via natural log, and VAT volume was transformed via cube root. Each variable included in the models was standardized to a mean of 0 and standard deviation of 1, and time was divided by 30 to correct scaling. Fasting and postprandial variables were standardized separately due to the inclusion of the fasting state as a covariate in the models. All data transformation and subsequent mixed-effects models were conducted in R (version 4.2.2) using the lmerTest^13^, performance^14^, and emmeans^15^ packages. For the interpretation of linear mixed-effects models, we assess significance using both the 95% confidence interval (e.g., does it cross 0) and the p-value (p < 0.05). Due to ambiguity in the determination of p-values for these models^16^, we deferred to the confidence interval when there was disagreement between the confidence intervals and p-values.

## RESULTS

There were 27 young adults (n=15 (56%) female) with no previously diagnosed metabolic pathology (**Table 1**). As expected, individuals with OW/OB had significantly (p < 0.001) higher body mass, volume of body fat, visceral fat area, and fat free mass as compared to individuals with NW. There were no significant differences in resting metabolic rate, RER, or substrate oxidation between individuals with OW/OB and NW (**Table 1**). Comparisons of participant characteristics based on biological sex are presented in **Supplementary Table 1**.

### Substrate Response to OGTT

Blood [glucose], serum [insulin], and plasma [lactate] significantly increased while plasma [NEFAs] significantly decreased following consumption of 100g glucose beverage (**Figure 1A-D**). Blood [glucose] (**Figure 1A**) was elevated above fasting levels (99.7 ± 8.5 mg/dL) at 30 (139.9 ± 21.0; p<0.001), 60 (124.9 ± 29.6; p<0.001), and 90 (115.3 ± 25.5; p = 0.007) minutes but not 120 minutes (109 ± 21.7; p = 0.214). [Glucose] area under the curve was not significantly different between individuals with obesity and individuals with normal weight BMI (15,076 ± 1912 mg/dl*min vs. 13,852 ± 2126 mg/d*minl; p = 0.13) Compared to fasting [insulin] (14.2 ± 8.1 µIU/mL), postprandial [insulin] was elevated at 30 (79.9 ± 75.1 µIU/mL; p < 0.001), 60 (77.9 ± 54.2 µIU/mL; p < 0.001), 90 (63.2 ± 47.7 µIU/mL; p = 0.006), and 120 (55.0 ± 36.4 µIU/mL; p = 0.03) minutes (all p<0.001; **Figure 1B**). [Insulin] area under the curve was not significantly different between individuals with obesity and individuals with normal weight BMI (9454 ± 5295 µIU/mL*min vs. 5929 ± 4284 µIU/mL*min; p = 0.09) Plasma [lactate] (**Figure 1C**) did not change from fasting (1.6 ± 0.4 mM) to 30 (1.5 ± 0.4 mM; p = 0.79), but significantly increased at 60 (1.9 ± 0.3 mM; P < 0.001), and 90 (1.8 ± 0.3 mM; p < 0.001) before returning to near baseline at 120 (1.7 ± 0.3; p = 0.06) minutes. [Lactate] area under the curve was not significantly different between individuals with obesity and individuals with normal weight BMI (205.7 ± 37.3 mM*minl vs. 208.9 ± 24.7 mM*min; p = 0.88) Plasma [NEFAs] (**Figure 1D**) were suppressed compared to fasting (0.41 ± 0.19 mEq/L) at 30 (0.28 ± 0.13 mEq/L), 60 (0.15 ± 0.08 mEq/L), 90 (0.15 ± 0.08 mEq/L), and 120 (0.09 ± 0.04 mEq/L) minutes (all p < 0.001). [NEFA] area under the curve was significantly higher for individuals with obesity than individuals with normal weight BMI (27.5 ± 10.4 mEq/L*min vs. 19.1 ± 7.2 mEq/L*min; p = 0.03).

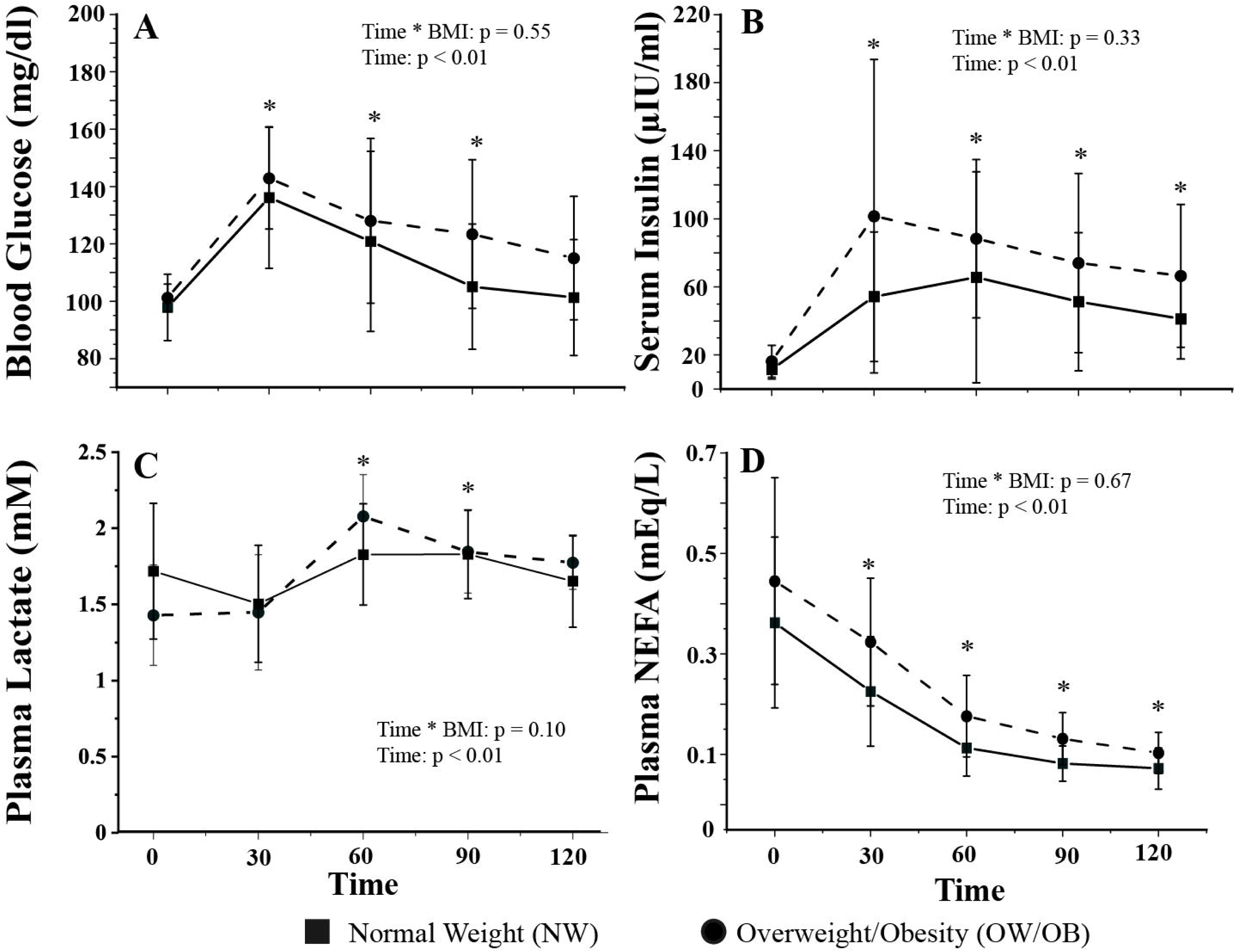

CHO (**Figure 2A**) was significantly elevated over time (time: p < 0.001), but there was no significant interaction between time and obesity status (time*BMI: p = 0.14). CHO was not different from fasting (0.102 ± 0.071 g/min) at 30 minutes (0.125 ± 0.071; p = 0.62) but was significantly higher at 60 (0.157 ± 0.063; p = 0.009), 90 (0.160 ± 0.049; p = 0.004), and 120 (0.166 ± 0.061; p = 0.002) minutes. There was a significant interaction between time and obesity status (time*BMI: p = 0.03) for FOX which indicated higher FOX throughout the postprandial period for individuals with OW/OB. Overall, FOX was unchanged from fasting (0.084 ± 0.024) at 30 (0.086 ± 0.024; p = 0.99) and 60 (0.078 ± 0.027; p = 0.83) minutes. Compared to fasting, FOX was suppressed at 90 (0.073 ± 0.026; p = 0.011) and 120 minutes (0.068 ± 0.023; p<0.001). RER (**Figure 2C**) was not different from fasting (0.795 ± 0.055) at 30 minutes (0.806 ± 0.052; p = 0.493) but was significantly higher at 60 (0.832 ± 0.051; p<0.001), 90 (0.839 ± 0.044; p<0.001), and 120 (0.845 ± 0.048; p<0.001) minutes. There was no significant time*BMI interaction (p = 0.08).

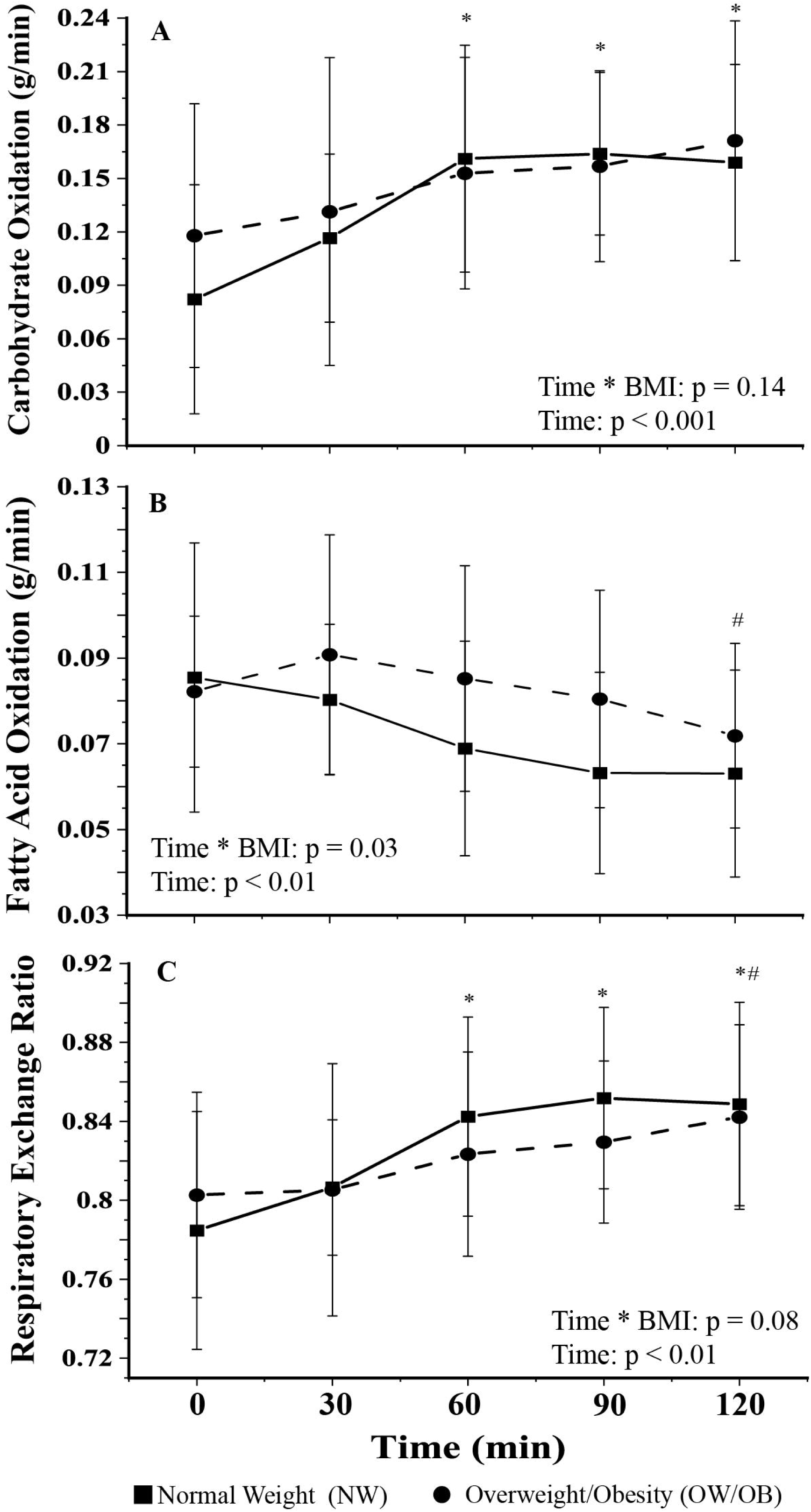

### Effect of body composition on postprandial metabolic flexibility (RER, CHO, and FOX)

We used mixed effects models to test for an effect of BF% on postprandial metabolic flexibility (**Table 4**). In these models, a quadratic model with a main effect for time^2^ and a time^2^ x BF% interaction term was a better fit to the postprandial data for RER (p = 0.002) and CHO (p = 0.002) than the linear model. In the quadratic models, BF% modified the relationship between postprandial RER and time (p = 0.011) and time^2^ (p = 0.019), as well as postprandial CHO and time (p = 0.014) and time^2^ (p = 0.023), indicating a nonlinear change in RER and CHO throughout the OGTT that depends on body composition. There was no evidence that BF% modifies the relationship between FOX and time (**Supplementary Table 2**). We also considered the effect of fat mass on postprandial RER and CHO. However, there was no evidence that fat mass modifies the relationship between postprandial RER and time (p = 0.364) or time^2^ (p = 0.409), as well as postprandial CHO and time (p = 0.364) or time^2^ (p = 0.409; see **Supplementary Table 3)**.

To assess the interaction of time and BF% and their effects on postprandial substrate oxidation as a measure of postprandial metabolic flexibility, we visualized the effect of time and BF% as continuous variables. We display (**Figure 3A-B**) the modeled RER and CHO for three representative values of BF%: the mean BF% of our study population, the mean BF% + 1 SD, and the mean BF% - 1 SD. The interaction of time and BF% displayed in these plots indicates that higher BF% is associated with higher postprandial RER (**Figure 3A**) and CHO (**Figure 3B**) demonstrating that higher BF% is associated with greater postprandial metabolic flexibility in our sample of young adults. The resulting visualization is adjusted for all covariates in **Table 4**.

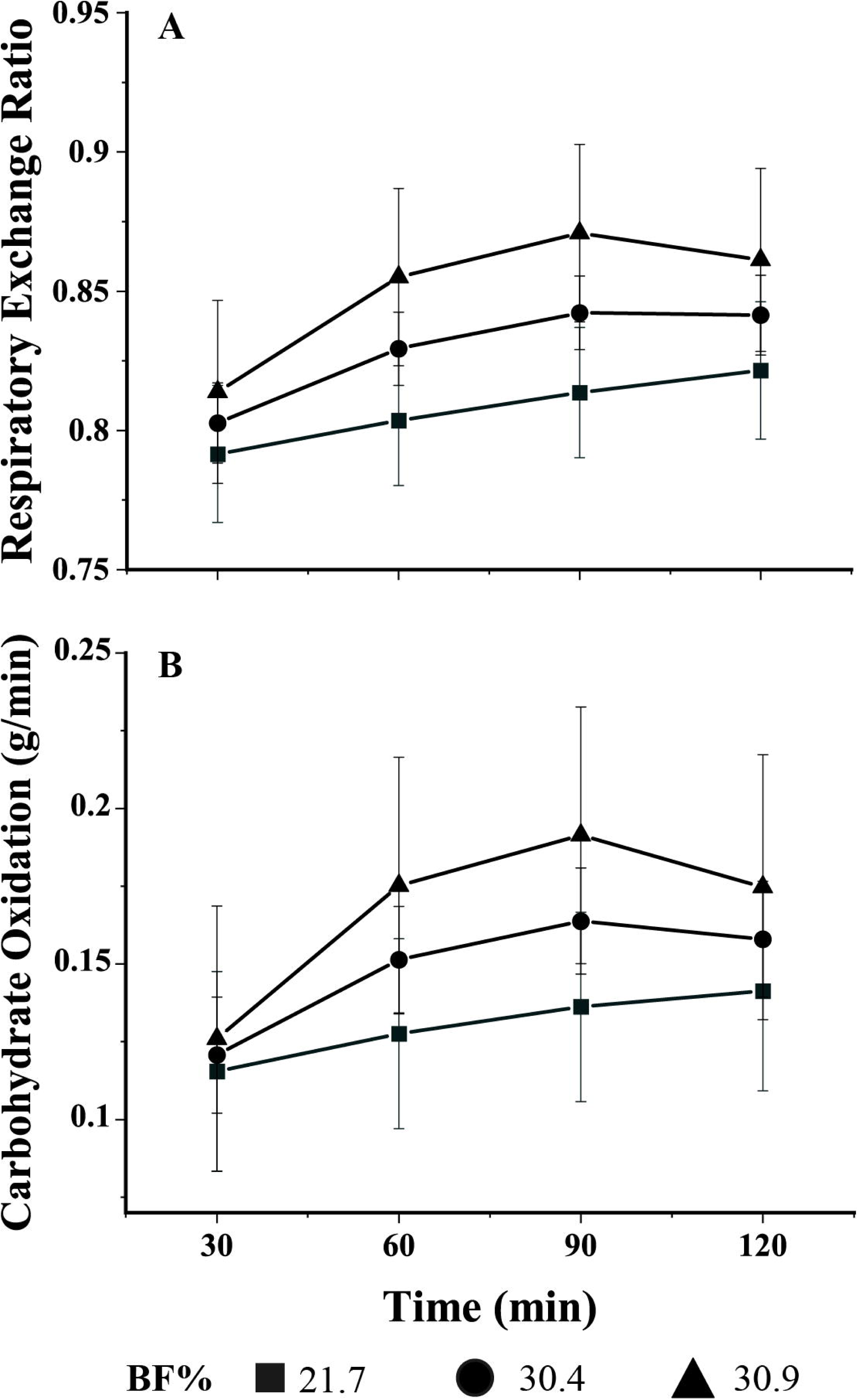

Further, greater insulin sensitivity (Matsuda Index) was associated with a higher postprandial RER or CHO (**Table 4**). Because skeletal muscle is a primary site of postprandial glucose disposal^17^ and people with obesity often have higher fat-free mass,^18^ we tested the effect of adding fat-free mass as an additional covariate. This did not improve the model fit (p = 0.204) over the original CHO model. Like the original CHO model, the addition of a time^2^ main effect and a time^2^ x BF% interaction term improved the model fit (p = 0.002), and percent body fat modified the relationship between CHO and time (p = 0.014) and time^2^ (p = 0.023) (**Table 4; Supplementary Results**). Like RER and CHO, higher BF% was associated with greater CHO adjusted for FFM, indicating differences in FFM across our sample do not affect the interaction between time and BF%.

## DISCUSSION

Metabolic flexibility is the ability to adjust substrate oxidation based on substrate availability or demand.^1^ The relationship between adiposity and measures of metabolic flexibility has been poorly analyzed and lacks clarity.^3, 19–24^ The majority of studies to date have categorized individuals based on BMI (an indicator of adiposity) and made inconsistent and inconclusive associations between adiposity and measures of metabolic flexibility.^3^ In this analysis, we analyzed the interaction between body fat percentage and postprandial metabolism (RER, CHO, and FOX) and demonstrate that higher body fat percentage is associated with elevated postprandial metabolic flexibility (RER and CHO) during an OGTT in young adults without overt metabolic pathology. These findings may seem paradoxical since individuals with higher BMI (e.g., obesity) are most often reported to have lower, or blunted, metabolic flexibility. However, this finding is not inconsistent with studies that show females have greater metabolic flexibility than males despite higher body fat^5^ and individuals with high metabolic flexibility had greater subcutaneous adipose tissue volumes.^7^ Furthermore, loss of subcutaneous adipose tissue during prolonged bed rest is accompanied by decreased metabolic flexibility.^20^ This study expands on previously published reports by directly analyzing the relationship between body fat and postprandial metabolic flexibility while adjusting for known covariates (i.e., age, sex, VAT, insulin sensitivity, etc.) that significantly affect substrate oxidation.

Adipose tissue function is proposed as a central regulator of metabolic flexibility^2^ by buffering fatty acid flux during the postprandial period.^4^ Obesity is associated with increased adipose tissue volume,^25^ elevated adipose tissue insulin resistance, adipose tissue inflammation, and altered adipokine release;^26^ all of which are associated with altered postprandial metabolism.^27^ In our sample of healthy young adults, higher body fat percentages did not alter suppression of lipolysis, as measured by plasma NEFAs, following ingestion of the glucose beverage. Furthermore, measures of adipose tissue insulin resistance (ADIP-IR) did not have a significant effect on postprandial RER or CHO. However, we cannot fully rule out reduced adipose tissue function in our findings. A critical role of adipose tissue in the postprandial state is glucose uptake and lipogenesis, which we cannot account for here.^28–30^ Importantly, our approach demonstrates that an increase in body fat percentage of ~6.9% (i.e., 20.0% to 26.9%) would result in an average increase of approximately 19.1 mg/dl of blood glucose at each time point during the OGTT (**Table 3**). This suggests the capacity for adipose tissue to buffer the blood glucose flux over time may be reduced in people with a higher body fat percentage, which would be consistent with observational studies in individuals with obesity, diabetes, and animal models.^31–34^ This function is an important part of adipose tissue’s role in reducing hyperglycemia in the postprandial period while subsequently storing nutrients for later oxidation.

**Table 3.** Effect of body composition on OGTT response.

| Characteristic<sup>1</sup> | Glucose |  |  | Insulin |  |  | NEFA |  |  |
| --- | --- | --- | --- | --- | --- | --- | --- | --- | --- |
|  | Beta<sup>2</sup> | 95% CI | p-value | Beta | 95% CI | p-value | Beta | 95% CI | p-value |
| Time | -0.55 | -0.723, -0.382 | **<0.001** | -0.17 | -0.286, -0.046 | **0.008** | -2.13 | -2.83, -1.44 | **<0.001** |
| Time<sup>2</sup> | --- | --- | --- | --- | --- | --- | 0.26 | 0.123, 0.397 | **<0.001** |
| BF% | 0.802 | 0.155, 1.45 | 0.055 | 0.232 | -0.175, 0.638 | 0.355 | 0.39 | -0.115, 0.894 | 0.216 |
| Fasting Measure | 0.231 | -0.061, 0.523 | 0.206 | -0.24 | -0.728, 0.239 | 0.412 | 0.211 | -0.067, 0.488 | 0.224 |
| Sex (Male) | 0.585 | -0.676, 1.84 | 0.451 | 0.058 | -0.681, 0.794 | 0.898 | 1.33 | 0.360, 2.31 | **0.030** |
| Age | -0.1 | -0.464, 0.267 | 0.66 | 0.016 | -0.190, 0.224 | 0.897 | 0.198 | -0.075, 0.472 | 0.244 |
| Matsuda | 0.188 | -0.247, 0.623 | 0.482 | -0.6 | -1.17, -0.018 | 0.105 | 0.08 | -0.242, 0.403 | 0.683 |
| AdipIR (OGTT) | 0.137 | -0.280, 0.554 | 0.591 | 0.438 | 0.205, 0.671 | **0.006** | 0.797 | 0.452, 1.14 | **0.001** |
| VAT volume<sup>1/3</sup> | -0.01 | -0.624, 0.595 | 0.969 | 0.056 | -0.285, 0.396 | 0.789 | -0.47 | -0.934, 0.001 | 0.116 |

<sup>1</sup>Fasting Measure denotes the fasting concentration of the outcome variable.
<sup>2</sup>A one-unit increase in time represents 30 minutes. A 1-SD increase in each variable corresponds to a 1-SD increase in glucose (19.1), insulin (0.59), or NEFA (0.45). Insulin and NEFA were transformed via natural log for normality. Variable SDs: BF%: 8.7; Fasting glucose: 8.5; Fasting insulin: 0.7; Fasting NEFA: 0.5; Age: 4.97; Matsuda: 2.9; AdipIR (OGTT): 13.3; VAT: 1.29.

**Table 4.** Effect of body composition (body fat percentage) on postprandial metabolic flexibility.

| Characteristic<sup>1</sup> | RER |  |  | CHO |  |  | CHO with FFM |  |  |
| --- | --- | --- | --- | --- | --- | --- | --- | --- | --- |
|  | Beta<sup>2</sup> | 95% CI | p-value | Beta | 95% CI | p-value | Beta | 95% CI | p-value |
| BF% x Time<sup>2</sup> | -0.133 | -0.240, -0.026 | **0.019** | -0.134 | -0.245, -0.023 | **0.023** | -0.134 | -0.245, -0.023 | **0.023** |
| BF% x Time | 0.732 | 0.189, 1.28 | **0.011** | 0.74 | 0.175, 1.31 | **0.014** | 0.74 | 0.175, 1.31 | **0.014** |
| BF% | -0.346 | -1.07, 0.374 | 0.39 | -0.51 | -1.26, 0.239 | 0.225 | -0.311 | -1.11, 0.492 | 0.502 |
| Time<sup>2</sup> | -0.156 | -0.263, -0.050 | **0.006** | -0.166 | -0.277, -0.056 | **0.005** | -0.166 | -0.277, -0.056 | **0.005** |
| Time | 1.08 | 0.537, 1.61 | **<0.001** | 1.06 | 0.498, 1.62 | **<0.001** | 1.06 | 0.498, 1.62 | **<0.001** |
| Fasting Measure | 0.803 | 0.549, 1.06 | **<0.001** | 0.793 | 0.474, 1.11 | **<0.001** | 0.7 | 0.358, 1.04 | **0.005** |
| Sex (Male) | 0.556 | -0.310, 1.42 | 0.3 | 0.421 | -0.487, 1.33 | 0.451 | 0.458 | -0.422, 1.34 | 0.414 |
| Age | -0.095 | -0.345, 0.155 | 0.536 | -0.07 | -0.332, 0.192 | 0.661 | -0.139 | -0.414, 0.137 | 0.429 |
| Matsuda Index | 0.505 | 0.207, 0.802 | **0.012** | 0.558 | 0.246, 0.869 | **0.009** | 0.59 | 0.285, 0.895 | **0.007** |
| AdipIR (OGTT) | 0.201 | -0.083, 0.486 | 0.255 | 0.123 | -0.176, 0.421 | 0.504 | 0.069 | -0.232, 0.370 | 0.715 |
| VAT volume<sup>1/3</sup> | -0.504 | -0.909, -0.098 | 0.055 | -0.224 | -0.648, 0.199 | 0.39 | -0.368 | -0.837, 0.100 | 0.224 |
| FFM | --- | --- | --- | --- | --- | --- | 0.305 | -0.177, 0.786 | 0.323 |

<sup>1</sup>Fasting Measure denotes the fasting measurement of the outcome variable. See main text for other abbreviations.
<sup>2</sup>A one-unit increase in time represents 30 minutes. A 1-SD increase in each covariate corresponds to a 1-SD increase in postprandial RER (0.044) or CHO (0.055 g/min). Covariate SDs: Fasting RER: 0.055; Fasting CHO: 0.071; FFM: 10.9; See Table 3 for other relevant SDs.

In the context of greater hyperglycemia following the OGTT, the elevated postprandial metabolic flexibility (postprandial RER and CHO) associated with higher body fat percentage in our sample would not be paradoxical. The postprandial period should induce an anabolic state (e.g. storage of nutrients) as compared to the catabolic state of fasting (e.g. breakdown of substrate).^4^ Metabolic flexibility is a critical metabolic adaptation to handle the complex integration of metabolism during caloric excess and restriction.^35^ Furthermore, it is essential for regular maintenance of blood glucose homeostasis following a feeding. Our observation of elevated CHO and RER in the setting of greater hyperglycemia suggests greater metabolic flexibility in people with higher body fat percentages, which would fit the concept of metabolic flexibility precisely; these individuals with higher body fat percentages are adjusting substrate oxidation based on the increased availability of glucose in the blood for longer periods of time. Individuals with lower body fat percentages who do not elevate CHO and RER during this period are likely experiencing the appropriate catabolic (storage) response to feeding by taking up and storing glucose as glycogen (liver and skeletal muscle) or fat (adipose tissue). As a result, they have no further need to increase CHO to maintain blood glucose homeostasis. A recent report by Curl et al.^36^ demonstrated altered glucose kinetics in older individuals as compared to younger adults without differences in metabolic flexibility, which they defined as the difference between peak and fasting RER. We did not assess glucose kinetics, but our findings on the impacts of body composition on postprandial blood glucose would be consistent with their findings.

This raises the question if our sample of young adults across the BMI spectrum but with no overt metabolic pathology are demonstrating a resilient, metabolically flexible phenotype.^37^ Metabolic flexibility is diminished following hypercaloric, high-fat feeding interventions and improved following significant weight loss, indicating it is a plastic phenotype.^38, 39^ No study has successfully tracked the disappearance of metabolic flexibility, but we suspect that long-term elevated levels of adiposity, such as that seen in chronic obesity, eventually lead to the loss of postprandial metabolic flexibility. In our sample, higher body fat percentage is inversely associated with insulin sensitivity (r =-0.616; p=0.001) while being positively associated with fasting values for serum insulin (r=0.427; p=0.037), plasma leptin (r=0.817; p<0.001), and adipose tissue insulin resistance (r=0.443; p=0.030). These relationships may be indicative or worsening metabolic health with higher body fat percentage, but the ability to elevate CHO oxidation during the OGTT (e.g. metabolic flexibility) demonstrates a resilient, metabolically flexibility phenotype in these individuals. The eventual loss of metabolic flexibility would therefore play a significant role in the progression of glucose dysfunction towards overt metabolic disease. That process and the mechanisms involved are still poorly understood and require both experimental and observational longitudinal studies.

Studies assessing the relationship between metabolic flexibility and adiposity primarily note inconsistent and inconclusive results which include positive associations, inverse associations, and no associations between indices of metabolic flexibility and measures of adiposity (e.g. total fat mas, visceral adipose volume).^3^ Some studies also rely on surrogates of adiposity such as BMI, waist circumference, or waist:hip ratio.^3, 22^ Though not the primary goal of our study, we note no significant differences in postprandial RER or CHO between individuals with normal weight and individuals with obesity as determined by the BMI; this is consistent with many studies. Directly analyzing the effect of body composition, while adjusting for covariates known to influence substrate oxidation, has allowed us to more clearly demonstrate an unexpected, but explainable, relationship in which higher body fat percentage is associated with postprandial metabolic flexibility in young adults without overt metabolic pathology. Assessing the relationship between body composition, instead of absolute fat mass or volume, and metabolic flexibility also accounts for the expected body mass differences between subjects and, more specifically, between biological sexes. By recruiting similarly from both sexes and including biological sex in our models, we account for potential sex differences independent of differences in body composition between the sexes. While our study is one of the larger studiesy analyzing metabolic flexibility to an OGTT in both males and females to date,^3^ we did not power our study to detect sex differences, which should be considered in future work. Recent reports demonstrate that total body mass as well as fat free mass are inversely associated with metabolic flexibility during an OGTT, highlighting the necessity of taking anthropometric differences into account.^40^

Furthermore, our approach to analyzing metabolic flexibility during an OGTT as a continuous variable across time is a novel aspect of our approach. Studies assessing metabolic flexibility during feeding challenges define metabolic flexibility as the change in RER or CHO from fasting to a defined timepoint (e.g. 60 min in an OGTT) or the peak change (max – min) in the measure.^3^ Our data, when analyzed in that way, produces similar results (**Supplementary Figure 1**). Defining postprandial metabolic flexibility based on a single timepoint during a dynamic and prolonged response, such as feeding, is likely a carryover from the EHC method, which applies indirect calorimetry to the steady-state phase of insulin infusion. Because postprandial metabolism is dynamic, a feeding challenge such as an OGTT has no equivalent to the steady-state period.^40, 41^ Curl et al.^36^ recently argued against postprandial RER as measure of metabolic flexibility following a glucose challenge when comparing younger vs. older adults, instead suggesting blood glucose kinetics as a better measure. We agree that the postprandial period should not be defined as a single measure and should be analyzed in a continuous manner across time. In fact, Curl et al.^36^ note significantly higher RER in their younger adults at 60-min postprandial but not at 120-min postprandial, highlighting the dynamic nature of this physiological process and the necessity to analyze it continuously.

We demonstrate that higher body fat percentage is associated with greater postprandial metabolic flexibility (RER and CHO) during an oral glucose tolerance test in young adults without overt metabolic pathology. We believe this finding is indicative of a preserved metabolically flexible phenotype where elevated postprandial blood glucose is resulting in elevated CHO with higher body fat percentage. Our approach of analyzing data in a continuous manner using linear mixed models, accounting for covariates known to modify the postprandial substrate oxidation response, adds clarity to the previously poorly defined relationship between adiposity and postprandial metabolic flexibility. While we cannot account for some functional (e.g. adipocyte size, lipogenesis, and molecular) or anatomical (e.g. distribution) aspects of adiposity, we more directly demonstrate what has long been ambiguous in published reports. These findings raise several new questions about adiposity and the continuum of metabolic health.

## Acknowledgements

The authors would like to acknowledge the Milken Institute School of Public Health Office of Research Excellence which provided funding for this project. The authors would also like to acknowledge the participants for their volunteered time and effort.

## Contributions

**NF:** Conceptualization, methodology, formal analysis, investigation, data curation, writing – original draft, visualization. **SR:** Investigation. **JB**: Investigation. **LS:** Investigation. **AL:** Investigation. **LD:** Conceptualization, writing – review & editing. **AC:** Formal analysis, writing – review & editing. **MB:** Conceptualization, methodology, investigation, writing – original draft, project administration, funding acquisition.

All authors have reviewed this manuscript and agree to its publication as written.

## References

1. Goodpaster BH, Sparks LM. Metabolic Flexibility in Health and Disease. Cell Metab. 2017;25(5):1027–36. doi: 10.1016/j.cmet.2017.04.015. PubMed PMID: 28467922; PMCID: PMC5513193.

2. Storlien L, Oakes ND, Kelley DE. Metabolic flexibility. Proc Nutr Soc. 2004;63(2):363–8. doi: 10.1079/PNS2004349. PubMed PMID: 15294056.

3. Glaves A, Diaz-Castro F, Farias J, Ramirez-Romero R, Galgani JE, Fernandez-Verdejo R. Association Between Adipose Tissue Characteristics and Metabolic Flexibility in Humans: A Systematic Review. Front Nutr. 2021;8:744187. Epub 20211203. doi: 10.3389/fnut.2021.744187. PubMed PMID: 34926544; PMCID: PMC8678067.

4. Frayn KN. Adipose tissue as a buffer for daily lipid flux. Diabetologia. 2002;45(9):1201–10. Epub 20020724. doi: 10.1007/s00125-002-0873-y. PubMed PMID: 12242452.

5. Sparks LM, Pasarica M, Sereda O, deJonge L, Thomas S, Loggins H, Xie H, Miles JM, Smith SR. Effect of adipose tissue on the sexual dimorphism in metabolic flexibility. Metabolism. 2009;58(11):1564–71. doi: 10.1016/j.metabol.2009.05.008. PubMed PMID: 19595383.

6. Sparks LM, Ukropcova B, Smith J, Pasarica M, Hymel D, Xie H, Bray GA, Miles JM, Smith SR. Relation of adipose tissue to metabolic flexibility. Diabetes Res Clin Pract. 2009;83(1):32–43. Epub 20081126. doi: 10.1016/j.diabres.2008.09.052. PubMed PMID: 19038471; PMCID: PMC2749984.

7. Fernandez-Verdejo R, Malo-Vintimilla L, Gutierrez-Pino J, Lopez-Fuenzalida A, Olmos P, Irarrazaval P, Galgani JE. Similar Metabolic Health in Overweight/Obese Individuals With Contrasting Metabolic Flexibility to an Oral Glucose Tolerance Test. Front Nutr. 2021;8:745907. Epub 20211116. doi: 10.3389/fnut.2021.745907. PubMed PMID: 34869522; PMCID: PMC8637191.

8. Taylor HL, Buskirk E, Henschel A. Maximal oxygen intake as an objective measure of cardio-respiratory performance. J Appl Physiol. 1955;8(1):73–80. doi: 10.1152/jappl.1955.8.1.73. PubMed PMID: 13242493.

9. Frayn KN. Calculation of substrate oxidation rates in vivo from gaseous exchange. J Appl Physiol Respir Environ Exerc Physiol. 1983;55(2):628–34. doi: 10.1152/jappl.1983.55.2.628. PubMed PMID: 6618956.

10. Matsuda M, DeFronzo RA. Insulin sensitivity indices obtained from oral glucose tolerance testing: comparison with the euglycemic insulin clamp. Diabetes Care. 1999;22(9):1462–70. doi: 10.2337/diacare.22.9.1462. PubMed PMID: 10480510.

11. Gastaldelli A, Gaggini M, DeFronzo RA. Role of Adipose Tissue Insulin Resistance in the Natural History of Type 2 Diabetes: Results From the San Antonio Metabolism Study. Diabetes. 2017;66(4):815–22. Epub 20170104. doi: 10.2337/db16-1167. PubMed PMID: 28052966.

12. Shashaj B, Luciano R, Contoli B, Morino GS, Spreghini MR, Rustico C, Sforza RW, Dallapiccola B, Manco M. Reference ranges of HOMA-IR in normal-weight and obese young Caucasians. Acta Diabetol. 2016;53(2):251–60. Epub 20150613. doi: 10.1007/s00592-015-0782-4. PubMed PMID: 26070771.

13. Kuznetsova A, Brockhoff, P.B., & Christensen R.H.B. lmerTest Package: Tests in Linear Mixed Effects Models. Journal of Statistical Software. 2017;82(13):1–26. doi: 10.18637/jss.v082.i13.

14. Lüdecke D, Ben-Shachar, M.S., Patil, I., Waggoner, P., Makowski, D. performance: An R Package for Assessment, Comparison and Testing of Statistical Models. The Journal of Open Source Software. 2021;6(60). doi: 10.21105/joss.03139.

15. Lenth R. emmeans: Estimated Margin2l Means, aka Least-Squares Means. R package version 1.10.3-090006. 2024.

16. Baayen RH, Davidson DJ, Bates DM. Mixed-effects modeling with crossed random effects for subjects and items. J Mem Lang. 2008;59(4):390–412. doi: 10.1016/j.jml.2007.12.005. PubMed PMID: WOS:000261651700003.

17. Kelley D, Mitrakou A, Marsh H, Schwenk F, Benn J, Sonnenberg G, Arcangeli M, Aoki T, Sorensen J, Berger M, et al. Skeletal muscle glycolysis, oxidation, and storage of an oral glucose load. J Clin Invest. 1988;81(5):1563–71. doi: 10.1172/JCI113489. PubMed PMID: 3130396; PMCID: PMC442590.

18. Dulloo AG, Jacquet J, Miles-Chan JL, Schutz Y. Passive and active roles of fat-free mass in the control of energy intake and body composition regulation. Eur J Clin Nutr. 2017;71(3):353–7. Epub 20161214. doi: 10.1038/ejcn.2016.256. PubMed PMID: 27966570.

19. Lightowler H, Schweitzer L, Theis S, Henry CJ. Changes in Weight and Substrate Oxidation in Overweight Adults Following Isomaltulose Intake During a 12-Week Weight Loss Intervention: A Randomized, Double-Blind, Controlled Trial. Nutrients. 2019;11(10). Epub 20191004. doi: 10.3390/nu11102367. PubMed PMID: 31590285; PMCID: PMC6836138.

20. Rudwill F, O’Gorman D, Lefai E, Chery I, Zahariev A, Normand S, Pagano AF, Chopard A, Damiot A, Laurens C, Hodson L, Canet-Soulas E, Heer M, Meuthen PF, Buehlmeier J, Baecker N, Meiller L, Gauquelin-Koch G, Blanc S, Simon C, Bergouignan A. Metabolic Inflexibility Is an Early Marker of Bed-Rest-Induced Glucose Intolerance Even When Fat Mass Is Stable. J Clin Endocrinol Metab. 2018;103(5):1910–20. doi: 10.1210/jc.2017-02267. PubMed PMID: 29546280; PMCID: PMC7263792.

21. Kahlhofer J, Lagerpusch M, Enderle J, Eggeling B, Braun W, Pape D, Muller MJ, Bosy-Westphal A. Carbohydrate intake and glycemic index affect substrate oxidation during a controlled weight cycle in healthy men. Eur J Clin Nutr. 2014;68(9):1060–6. Epub 20140709. doi: 10.1038/ejcn.2014.132. PubMed PMID: 25005676.

22. Bergouignan A, Antoun E, Momken I, Schoeller DA, Gauquelin-Koch G, Simon C, Blanc S. Effect of contrasted levels of habitual physical activity on metabolic flexibility. J Appl Physiol (1985). 2013;114(3):371–9. Epub 20121213. doi: 10.1152/japplphysiol.00458.2012. PubMed PMID: 23239872.

23. Assaad M, El Mallah C, Obeid O. Phosphorus ingestion with a high-carbohydrate meal increased the postprandial energy expenditure of obese and lean individuals. Nutrition. 2019;57:59–62. Epub 20180619. doi: 10.1016/j.nut.2018.05.019. PubMed PMID: 30153580.

24. Huda MS, Dovey T, Wong SP, English PJ, Halford J, McCulloch P, Cleator J, Martin B, Cashen J, Hayden K, Wilding JP, Pinkney J. Ghrelin restores ‘lean-type’ hunger and energy expenditure profiles in morbidly obese subjects but has no effect on postgastrectomy subjects. Int J Obes (Lond). 2009;33(3):317–25. Epub 20090203. doi: 10.1038/ijo.2008.270. PubMed PMID: 19188925.

25. Turer AT, Khera A, Ayers CR, Turer CB, Grundy SM, Vega GL, Scherer PE. Adipose tissue mass and location affect circulating adiponectin levels. Diabetologia. 2011;54(10):2515–24. Epub 20110722. doi: 10.1007/s00125-011-2252-z. PubMed PMID: 21779869; PMCID: PMC4090928.

26. Shafiei MS, Shetty S, Scherer PE, Rockey DC. Adiponectin regulation of stellate cell activation via PPARgamma-dependent and -independent mechanisms. Am J Pathol. 2011;178(6):2690–9. doi: 10.1016/j.ajpath.2011.02.035. PubMed PMID: 21641391; PMCID: PMC3124230.

27. Sun K, Kusminski CM, Scherer PE. Adipose tissue remodeling and obesity. J Clin Invest. 2011;121(6):2094–101. Epub 20110601. doi: 10.1172/JCI45887. PubMed PMID: 21633177; PMCID: PMC3104761.

28. Chadt A, Al-Hasani H. Glucose transporters in adipose tissue, liver, and skeletal muscle in metabolic health and disease. Pflugers Arch. 2020;472(9):1273–98. Epub 20200626. doi: 10.1007/s00424-020-02417-x. PubMed PMID: 32591906; PMCID: PMC7462924.

29. Luo L, Liu M. Adipose tissue in control of metabolism. J Endocrinol. 2016;231(3):R77–R99. doi: 10.1530/JOE-16-0211. PubMed PMID: 27935822; PMCID: PMC7928204.

30. Roberts R, Hodson L, Dennis AL, Neville MJ, Humphreys SM, Harnden KE, Micklem KJ, Frayn KN. Markers of de novo lipogenesis in adipose tissue: associations with small adipocytes and insulin sensitivity in humans. Diabetologia. 2009;52(5):882–90. Epub 20090228. doi: 10.1007/s00125-009-1300-4. PubMed PMID: 19252892.

31. Fabbrini E, Yoshino J, Yoshino M, Magkos F, Tiemann Luecking C, Samovski D, Fraterrigo G, Okunade AL, Patterson BW, Klein S. Metabolically normal obese people are protected from adverse effects following weight gain. J Clin Invest. 2015;125(2):787–95. Epub 20150102. doi: 10.1172/JCI78425. PubMed PMID: 25555214; PMCID: PMC4319438.

32. Graham TE, Kahn BB. Tissue-specific alterations of glucose transport and molecular mechanisms of intertissue communication in obesity and type 2 diabetes. Horm Metab Res. 2007;39(10):717–21. doi: 10.1055/s-2007-985879. PubMed PMID: 17952832.

33. Herman MA, Peroni OD, Villoria J, Schon MR, Abumrad NA, Bluher M, Klein S, Kahn BB. A novel ChREBP isoform in adipose tissue regulates systemic glucose metabolism. Nature. 2012;484(7394):333–8. doi: 10.1038/nature10986. PubMed PMID: 22466288; PMCID: PMC3341994.

34. Hoffstedt J, Forster D, Lofgren P. Impaired subcutaneous adipocyte lipogenesis is associated with systemic insulin resistance and increased apolipoprotein B/AI ratio in men and women. J Intern Med. 2007;262(1):131–9. doi: 10.1111/j.1365-2796.2007.01811.x. PubMed PMID: 17598821.

35. Smith RL, Soeters MR, Wust RCI, Houtkooper RH. Metabolic Flexibility as an Adaptation to Energy Resources and Requirements in Health and Disease. Endocr Rev. 2018;39(4):489–517. doi: 10.1210/er.2017-00211. PubMed PMID: 29697773; PMCID: PMC6093334.

36. Curl CC, Leija RG, Arevalo JA, Osmond AD, Duong JJ, Huie MJ, Masharani U, Horning MA, Brooks GA. Altered glucose kinetics occurs with aging: a new outlook on metabolic flexibility. Am J Physiol Endocrinol Metab. 2024;327(2):E217–E28. Epub 20240619. doi: 10.1152/ajpendo.00091.2024. PubMed PMID: 38895979; PMCID: PMC11427093.

37. Mietus-Snyder M, Perak AM, Cheng S, Hayman LL, Haynes N, Meikle PJ, Shah SH, Suglia SF, American Heart Association Atherosclerosis H, Obesity in the Young Committee of the Council on Lifelong Congenital Heart D, Heart Health in the Y, Council on L, Cardiometabolic H, Council on Arteriosclerosis T, Vascular B, Council on C, Stroke N. Next Generation, Modifiable Cardiometabolic Biomarkers: Mitochondrial Adaptation and Metabolic Resilience: A Scientific Statement From the American Heart Association. Circulation. 2023;148(22):1827–45. Epub 20231030. doi: 10.1161/CIR.0000000000001185. PubMed PMID: 37902008.

38. Berk ES, Kovera AJ, Boozer CN, Pi-Sunyer FX, Albu JB. Metabolic inflexibility in substrate use is present in African-American but not Caucasian healthy, premenopausal, nondiabetic women. J Clin Endocrinol Metab. 2006;91(10):4099–106. Epub 20060725. doi: 10.1210/jc.2005-2411. PubMed PMID: 16868062; PMCID: PMC2670464.

39. Kelley DE, Goodpaster B, Wing RR, Simoneau JA. Skeletal muscle fatty acid metabolism in association with insulin resistance, obesity, and weight loss. Am J Physiol. 1999;277(6):E1130–41. doi: 10.1152/ajpendo.1999.277.6.E1130. PubMed PMID: 10600804.

40. Alcantara JMA, Galgani JE. Association of metabolic flexibility indexes after an oral glucose tolerance test with cardiometabolic risk factors. Eur J Clin Nutr. 2024;78(3):180–6. Epub 20231218. doi: 10.1038/s41430-023-01373-w. PubMed PMID: 38110728.

41. Alcantara JMA, Sanchez-Delgado G, Jurado-Fasoli L, Galgani JE, Labayen I, Ruiz JR. Reproducibility of the energy metabolism response to an oral glucose tolerance test: influence of a postcalorimetric correction procedure. Eur J Nutr. 2023;62(1):351–61. Epub 20220825. doi: 10.1007/s00394-022-02986-w. PubMed PMID: 36006468; PMCID: PMC9899729.

