## Supplementary for "Increased Postprandial Metabolic Flexibility is Associated with Higher Body Fat Percentages in Healthy Young Adults"

**Supplemental Materials**

*Effect of body composition on substrate response*

Using mixed-effects models, we tested the effect of BF% on glucose, insulin, and NEFAs over time and time^2^ to account for a potential nonlinear response. To interpret our results, obtained beta values were multiplied by 8.7%, the standard deviation for body fat percentage in our sample, to demonstrate the expected changes in blood glucose, serum insulin, and serum NEFAs based on the standard deviations for those measures (**Table 3**). The addition of a time^2^ main effect and a time^2^ x BF% interaction did not improve the model fit for the effect of BF% on postprandial blood glucose (p = 0.237) or insulin (p = 0.545), suggesting that a linear model has adequate fit relative to a quadratic model. In the linear models without time^2^, BF% did not modify the relationship between time and glucose (p = 0.294) or insulin (p = 0.178). In the subsequent main effects models, a 6.9% increase in body fat percentage would result in a significant (β = 0.802; 95% CI: 0.155, 1.45; p = 0.055) 19.1 mg/dl increase in postprandial blood glucose at each time point. A 2.0% change in body fat is associated with a non-significant increase in serum insulin (β = 0.232; 95% CI: -0.175, 0.638; p = 0.355). For NEFAs, model fit was improved by the addition of a time^2^ main effect and a time^2^ x BF% interaction (p < 0.001) compared to the model with only linear time terms. In this quadratic model, BF% did not modify the relationship between NEFAs and time (p = 0.082) or time^2^ (p = 0.313). In the main effects model without the interaction term, a 3.4% body fat percent increase resulted in a non-significant (β = 0.39; 95% CI: -0.115, 0.894; p = 0.216) increase postprandial plasma NEFA 1.19 (mEq/L). Thus, body composition affects postprandial blood glucose but not serum insulin or NEFAs throughout the OGTT (**Table 3**).

| Supplementary Table 1. Participant Anthropometric Characteristics based on biological sex | | | | |
| --- | --- | --- | --- | --- |
| **Variable** | **All (n = 27)** | **Female (n = 15)** | **Male (n = 12)** | **p-value (Sex)** |
| NIH Race/Ethnicity (%) |  |  |  | 0.561 |
| Black | 7 (25.9%) | 4 (26.7%) | 3 (25.0%) | --- |
| Hispanic or Latino | 1 (3.7%) | 0 (0.0%) | 1 (8.3%) | --- |
| Unknown | 1 (3.7%) | 1 (6.7%) | 0 (0.0%) | --- |
| White | 18 (66.7%) | 10 (66.7%) | 8 (66.7%) | --- |
| Age (years) | 22 [19, 27] | 22 [19, 24] | 25 [21, 28] | 0.138 |
| Height (m) | 1.7 (0.09) | 1.6 (0.05) | 1.8 (0.05) | **<0.001** |
| Weight (kg) | 73.0 [66.6, 90.0] | 72.4 [63.5, 87.7] | 76.8 [71.1, 100.0] | 0.075 |
| BMI (kg/m^2^) | 27.1 (4.5) | 27.7 (4.7) | 26.4 (4.4) | 0.474 |
| Fat Mass (kg) | 24.2 (9.6) | 26.3 (8.6) | 21.7 (10.6) | 0.221 |
| Fat Free Mass (kg) | 51.7 (10.9) | 45.2 (6.3) | 59.7 (10) | **<0.001** |
| Body Fat (%) | 30.4 (8.7) | 34.9 (6.3) | 24.7 (8.1) | **0.001** |
| VAT Volume (in^3^) | 2.7 (1.3) | 2.2 (1.3) | 3.3 (1.0) | **0.035** |
| Resting Metabolic Rate (kcal/d) | 1709.8 (287.9) | 1538.2 (222.3) | 1924.2 (206.8) | **<0.001** |
| Respiratory Exchange Ratio (RER) | 0.80 (0.06) | 0.78 (0.05) | 0.82 (0.06) | 0.056 |
| Carbohydrate Oxidation (g/min) | 0.1 (0.07) | 0.07 (0.04) | 0.14 (0.08) | **0.005** |
| Lipid Oxidation (g/min) | 0.08 (0.02) | 0.08 (0.03) | 0.08 (0.02) | 0.91 |
| VO_2PEAK_ (mL/kg/min) | 34.1 (8.8) | 31.4 (7.3) | 37.5 (9.7) | 0.075 |
| VO_2PEAK_ (L/min) | 2.6 (0.6) | 2.3 (0.37) | 3.1 (0.53) | **<0.001** |
| Race/ethnicity is reported as count (percent). Numerical data are mean ± SD or median [IQR] based on normality. | | | | |

| **Supplementary Table 2.** Effect of BF% on Postprandial Fat Oxidation (FOX) as a Measure of Metabolic Flexibility | | | |
| --- | --- | --- | --- |
| **Characteristic** | **Beta** | **95% CI** | **p-value** |
| Time | -0.27 | -0.356, -0.184 | **<0.001** |
| BF% | -0.334 | -0.754, 0.086 | 0.203 |
| Fasting Measure | 0.689 | 0.469, 0.908 | **<0.001** |
| Sex (Male) | -0.4 | -1.23, 0.426 | 0.431 |
| Age | 0.135 | -0.111, 0.382 | 0.374 |
| Matsuda | -0.394 | -0.689, -0.099 | **0.041** |
| AdipIR (OGTT) | -0.217 | -0.490, 0.056 | 0.204 |
| VAT volume^1/3^ | 0.457 | 0.054, 0.861 | 0.077 |

| **Supplementary Table 3.** Effect of Fat Mass on Measures of Postprandial Metabolic Flexibility | | | | | | | | | |
| --- | --- | --- | --- | --- | --- | --- | --- | --- | --- |
|  | RER | | | CHO | | | CHO with FFM | | |
| **Characteristic** | **Beta** | **95% CI** | **p-value** | **Beta** | **95% CI** | **p-value** | **Beta** | **95% CI** | **p-value** |
| Time^2^ | -0.147 | -0.259, -0.036 | **0.012** | -0.157 | -0.273, -0.041 | **0.01** | -0.157 | -0.273, -0.041 | **0.01** |
| Time | 1.03 | 0.459, 1.59 | **<0.001** | 1.01 | 0.419, 1.60 | **0.001** | 1.01 | 0.419, 1.60 | **0.001** |
| Fat Mass | 0.367 | -0.160, 0.894 | 0.263 | 0.361 | -0.157, 0.880 | 0.263 | 0.362 | -0.156, 0.881 | 0.277 |
| Fasting Measure | 0.761 | 0.489, 1.03 | **<0.001** | 0.732 | 0.409, 1.06 | **0.002** | 0.721 | 0.367, 1.07 | **0.005** |
| Sex (Male) | 0.194 | -0.610, 0.999 | 0.693 | 0.309 | -0.518, 1.14 | 0.542 | 0.283 | -0.608, 1.17 | 0.616 |
| Age | -0.148 | -0.413, 0.116 | 0.365 | -0.107 | -0.371, 0.156 | 0.506 | -0.116 | -0.400, 0.168 | 0.52 |
| Matsuda | 0.449 | 0.136, 0.762 | **0.03** | 0.544 | 0.236, 0.852 | **0.01** | 0.541 | 0.231, 0.851 | **0.013** |
| AdipIR (OGTT) | 0.145 | -0.154, 0.444 | 0.43 | 0.074 | -0.223, 0.372 | 0.683 | 0.066 | -0.247, 0.380 | 0.737 |
| VAT volume^1/3^ | -0.468 | -0.963, 0.028 | 0.135 | -0.268 | -0.750, 0.213 | 0.367 | -0.271 | -0.755, 0.212 | 0.379 |
| Fat-free Mass | --- | --- | --- | --- | --- | --- | 0.033 | -0.373, 0.438 | 0.899 |

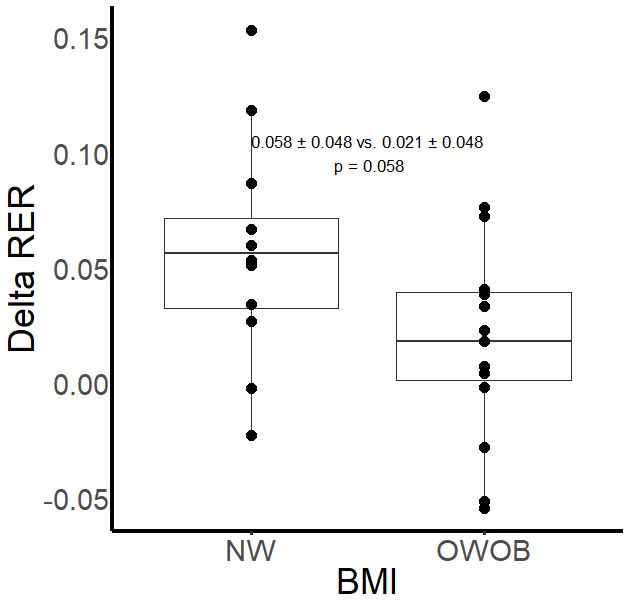

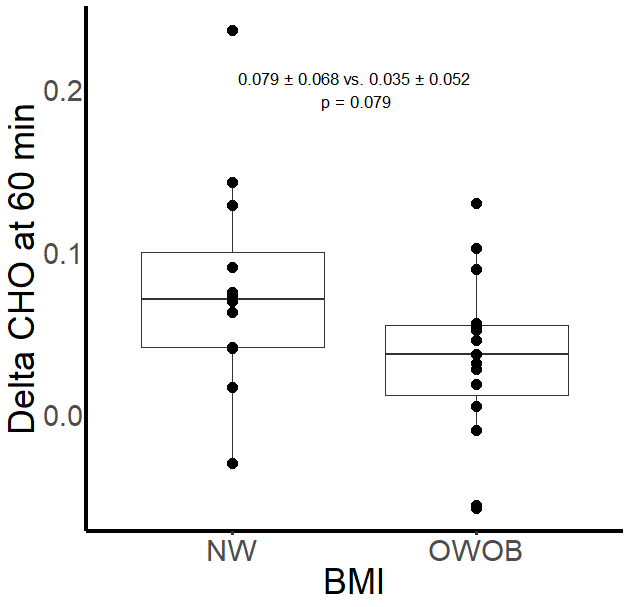

**Supplementary Figure 1.** Metabolic flexibility and individuals with normal weight (NW) or Overweight and Obesity (OWOB) as determined by Delta RER and Delta CHO. Differences between groups (BMI classification) was compared via T-test.
